# LSP1-myosin1e bi-molecular complex regulates focal adhesion dynamics and cell migration

**DOI:** 10.1101/2020.02.26.963991

**Authors:** Katja Schäringer, Sebastian Maxeiner, Carmen Schalla, Stephan Rütten, Martin Zenke, Antonio Sechi

**Author notes:** equal contribution (Katja Schäringer, Sebastian Maxeiner).

## Abstract

Several cytoskeleton-associated proteins and signalling pathways work in concert to regulate actin cytoskeleton remodelling, cell adhesion and migration. We have recently demonstrated that the bi-molecular complex between the leukocyte-specific protein 1 (LSP1) and myosin1e controls actin cytoskeleton remodelling during phagocytosis. In this study, we show that LSP1 down regulation severely impairs cell migration, lamellipodia formation and focal adhesion dynamics in macrophages. Inhibition of the interaction between LSP1 and myosin1e also impairs these processes resulting in poorly motile cells, which are characterised by few and small lamellipodia. Furthermore, cells in which LSP1-myosin1e interaction is inhibited are typically associated with inefficient focal adhesion turnover. Collectively, our findings show that the LSP1-myosin1e bimolecular complex plays a pivotal role in the regulation of actin cytoskeleton remodelling and focal adhesion dynamics required for cell migration.

## Introduction

A large library of actin-associated proteins steer nucleation, cross-linking, capping and elongation of actin filaments. The precise spatial and temporal co-ordination of these functions is fundamental for the movement of cells that is required for many biological events ranging from organ development to tissue repair. The importance of actin cytoskeleton dynamics is emphasised by the onset and progress of diseases due to cells lacking or expressing mutated variants of actin-associated proteins (Mathieson, 2012; Ramaekers and Bosman, 2004). In spite of several studies, the functions of some actin-associated proteins have not been well defined. One of such proteins is the leukocyte-specific protein 1 (LSP1). LSP1 is expressed in several cell types of the immune system such as T-cells, B-cells, macrophages and neutrophils. It is also expressed in myeloid and lymphoid cell lines and, despite its name, in endothelial cells (Jongstra et al., 1994; Jongstra et al., 1988; Jongstra-Bilen et al., 2000; Kadiyala et al., 1990; Liu et al., 2005; Maxeiner et al., 2015; Palker et al., 1998).

The amino-terminal half of LSP1 incorporates Ca^2+^-binding sites and a coiled-coil region (Jongstra et al., 1988; Klein et al., 1989), suggesting that Ca^2+^ signalling and dimerization could regulate LSP1 function. The carboxy-terminal half incorporates a caldesmon-like region having a weaker F-actin-binding activity (Zhang et al., 2000; Zhang et al., 2001) and two villin headpiece-like sequences, which primarily mediate the interaction of LSP1 with F-actin (Klein et al., 1990; Wong et al., 2003; Zhang et al., 2001). We have demonstrated that the carboxy-terminal half of LSP1 directly interacts with the SH3 domain of the molecular motor myosin1e through the non-canonical SH3-binding site AGDMSKKS (Maxeiner et al., 2015). These studies suggest that LSP1 may be involved in the regulation of actin cytoskeleton architecture and dynamics. Indeed, the actin-binding activity of LSP1 is required for the formation of the long, actin-rich cell projections that develop in a wide-ranging variety of cells, which overexpress LSP1 (Howard et al., 1998; Miyoshi et al., 2001; Zhang et al., 2001).

We have provided a direct evidence that LSP1 regulates actin cytoskeleton dynamics. We found that LSP1 localisation and dynamics at internalisation sites during Fcγ receptor-mediated phagocytosis, a process that depends on actin dynamics, spatially and temporally overlap with that of the actin cytoskeleton (Maxeiner et al., 2015). Moreover, in LSP1-deficient macrophages and in macrophages in which LSP1-myosin1e or LSP1-actin interactions are inhibited, Fcγ receptor-mediated phagocytosis is severely reduced (Maxeiner et al., 2015). Given the modulation of actin dynamics by LSP1, it is not surprising that LSP1 has been implicated in the regulation of migration of several cell types including neutrophils, dendritic cells and T-cells (Coates et al., 1991; Howard et al., 1994; Howard et al., 1998; Hwang et al., 2015; Jongstra-Bilen et al., 2000; Koral et al., 2015; Li et al., 2000; Petri et al., 2011).

Although these studies clearly show that LSP1 is involved in the regulation of actin cytoskeleton structural organisation and dynamics, the molecular mechanisms underlying the function of this actin-associated protein are still poorly characterised. Current evidence shows that LSP1 is phosphorylated at serine and threonine sites (Carballo et al., 1996; Huang et al., 1997; Jongstra-Bilen et al., 1990; Matsumoto et al., 1995a; Matsumoto et al., 1993; Wu et al., 2007). In lymphocytes, LSP1 is phosphorylated by protein kinase C (PKC) (Carballo et al., 1996; Matsumoto et al., 1995b; Matsumoto et al., 1993), whereas in neutrophils stimulated with the chemoattractant formyl-methionyl-leucyl-phenylalanine (fMLP), LSP1 is phosphorylated by the mitogen-activated protein (MAP) kinase–activated protein kinase 2 (MK2) (Huang et al., 1997; Wu et al., 2007). Notably, PKC-dependent phosphorylation of LSP1 decreases its localisation with the plasma membrane and the actin cytoskeleton (Matsumoto et al., 1995b; Miyoshi et al., 2001). By contrast, LSP1 phosphorylated by MK2 results in the accumulation of phosphorylated LSP1 at the leading edge of neutrophils (Wu et al., 2007). The importance of the interaction between kinases and LSP1 is further supported by the observation that LSP1 targets proteins of the ERK/MAP kinase pathway to the actin cytoskeleton (Harrison et al., 2004). Thus, it is plausible that the localisation of LSP1 to actin-rich structures depends on its phosphorylation status and can be regulated by diverse kinases and signalling pathways.

Regardless the positive or negative regulation of cell migration, it is unquestionable that LSP1 controls this important biological process. By contrast, very little is known about the molecular mechanisms underlying this LSP1 function. For instance, it has been shown that LSP1 participates in a complex with WASP and the Arp2/3 complex (Prasad et al., 2012), two important regulators of actin filament nucleation. Furthermore, LSP1 can also be found in a complex together myosin IIA and one of its regulators, the myosin light chain kinase (Cervero et al., 2018). Since LSP1 does not directly interact with WASP, the Arp2/3 complex and myosin IIA, it is likely that LSP1 is recruited to these complexes via its interaction with F-actin. Notably, we have demonstrated that LSP1 binds to the SH3 domain of myosin1e and that this bi-molecular complex is essential for efficient actin cytoskeleton dynamics during Fcγ receptor-mediated phagocytosis (Maxeiner et al., 2015).

In this study, we have added another piece to the puzzle describing the *modus operandi* of LSP1. We have demonstrated that the interaction of LSP1 with myosin 1e is essential for efficient focal adhesion dynamics and zyxin kinetics at these locations. The LSP1-myosin1e binary complex also regulates lamellipodia formation and dynamics. Consequently, interfering with LSP1-myosin1e interaction impaired cell migration.

## Materials and Methods

### Cell culture

Wild type and genetically modified J774 macrophage cell lines were grown in DMEM supplemented with 10% fetal calf serum (FCS), 4 mM L-glutamine, 100 μg/mL streptomycin, and 100 U/mL penicillin. The packaging cell line 293T (CRL 11268; ATCC) was grown in DMEM high glucose supplemented with 10% FCS, 2 mM L-glutamine, 1 mM sodium pyruvate, 100 μg/mL streptomycin, and 100 U/mL penicillin. All cell lines were grown at 37°C and 5% CO_2_.

### Cloning and generation of genetically modified J774 cells

To generate RFP-zyxin, the coding sequence of zyxin was amplified with the following primer pair: forward 5’-GCTTCGAATTCCATGGCGGCCCCCCGCCCGTCT-3’ (containing a EcoRI site) and reverse 5’-CTCGAGGATCCTCAGGTCTGGGCTCTAGCAGTGTGGCA-3’ (containing a BamHI site) using pMSCV-RFP-Zyxin as the template (Gamper et al., 2016). The amplified product was then cloned into the EcoRI and BamHI site of pWPXL-RFP (Maxeiner et al., 2015). Turquoise-zyxin was cloned as following. The coding sequence of turquoise was amplified from pLL3.7m-mTurquoise2-SLBP (18-126)-IRES-H1-mMaroon1 (Addgene vector no. 83842) using the following primer pair: forward 5’-CGTTTAAACAGGTATGGTGAGCAAGGGCGA-3’ (containing a PmeI site) and reverse 5’-GCAGCGAATTCCCTCCCAGGGAACGCAACATTGAGTA-3’ (containing an EcoRI site). The amplified product was then cloned into pWPXL-RFP-Zyxin after excision of RFP using PmeI and EcoRI to generate pWPXL-Turquoise-zyxin. Both RFP-zyxin and turquoise-zyxin were sequenced to verify the accuracy of the cloning procedure. To generate cells expressing RFP-zyxin or turquoise-zyxin, wild-type and LSP1-KD J774 cells were transduced with lentiviruses carrying the RFP-zyxin gene, whereas J774 cells expressing the deletion mutant LSP1-ΔSBS or full-length LSP1 were transduced with lentiviruses carrying the turquoise-zyxin gene. Generation of lentiviruses and transduction of cells was done as already described (Maxeiner et al., 2015).

### Immunofluorescence and live cell imaging

Immunofluorescence labelling was done as previously described (Gamper et al., 2016; Maxeiner et al., 2015; Sechi et al., 2016). For vinculin labelling, cells were fixed with 1% paraformaldehyde (PFA)/0.5% Triton X-100 in cytoskeleton buffer for 15 min at RT and then post-fixed with 4% PFA in cytoskeleton buffer for 10 min at RT. For EB1 and tubulin labelling, cells were fixed with ice-cold (−20°C) methanol for 4 min, rehydrated with 0.1% Triton X-100 in Tris-buffered saline (TBS; 3x, 5 min), and finally washed with TBS. Vinculin, EB1 and tubulin were detected with the monoclonal antibody hVin1 (Sigma-Aldrich), clone 5 (BD Transduction Laboratories, Heidelberg, Germany) and the rat hybridoma supernatant YL1/2 (Wehland et al., 1983), respectively. The actin cytoskeleton was visualised with Alexa fluorophore-conjugated phalloidin (Life Technologies). For live cell imaging, phase contrast and epifluorescence images were acquired with an Axio Observer Z1 inverted microscope (Carl Zeiss, Jena, Germany) equipped with an EMCCD camera (Evolve Delta, Photometrics, Tucson, AZ) driven by ZEN 2.3 software (Carl Zeiss, Jena, Germany).

### Analysis of cell migration and focal adhesion dynamics

J774 cells were plated onto self-made glass-bottomed dishes (ø 6 cm) and their migration was recorded continuously for 24 h (images were acquired every 5 min). The migration of all J774 cell lines was analysed using the Fiji (https://imagej.net/Fiji) plug-in MTrackJ (Meijering et al., 2012) to quantify parameters such as average speed and directionality. Cells that touched neighbouring cells, diving cells and cells that displayed an oscillating movement were excluded from the analysis. Focal adhesion dynamics was analysed as previously described (Berginski and Gomez, 2013; Würflinger et al., 2011).

### Analysis of lamellipodia dynamics

Lamellipodia dynamics was visualised by phase contrast microscopy after plating J774 cells at low density onto self-made glass-bottomed dishes (ø 6 cm). Phase contrast images were acquired every 5 seconds using an EMCCD camera (Cascade 512B, Photometrics, Tucson, AZ, USA) driven by IPLab Spectrum software (Scanalytics, Fairfax, VA, USA). The following parameters were measured: number of cells associated with lamellipodia (% of total cell number), velocity of lamellipodia spreading and lamellipodia width (measured from the beginning of lamellipodia spreading until the first signs of lamellipodia retraction).

### Total internal reflection fluorescence microscopy (TIRF)

TIRF microscopy was performed on an Axio Observer Z1 inverted microscope equipped with a motorized TIRF slider (Zeiss). Excitation of GFP, RFP and Turquoise was done using 488, 561 and 458 nm laser lines (at 10% of their nominal output power for 488 and 561, 30% for 458), respectively. The depth of the evanescent field for all wavelengths was ∼70 nm. Images were acquired every 10 seconds using an Evolve Delta EMCCD camera driven by ZEN software (Zeiss). For all experiments, exposure time, depth of the evanescent field, and electronic gain were kept constant.

### Fluorescence recovery after photobleaching (FRAP)

To analyse focal adhesion kinetics, J774 cells expressing RFP- or Turquoise-tagged zyxin were seeded onto self-made glass-bottomed dishes (ø 6 cm). For fluorescence recovery after photobleaching, cells were imaged on an Axio Observer Z1 inverted microscope equipped with heating stage and CO_2_ controller (Zeiss) maintained at a constant temperature of 37°C. A portion of single focal adhesions (approximately ø 3.84 µm) was photobleached using a 405 nm laser driven by the UGA-40 control unit (Rapp Opto Electronic GmbH, Wedel, Germany). The recovery of the fluorescent signal was monitored by imaging cells every second for 15 min. Imaging was done using an Evolve Delta EMCCD camera driven by ZEN software (Zeiss). For all experiments the size of the bleached area, and the duration and intensity of the laser impulse were kept constant. The extent of recovery of the fluorescent signal was determined using Fiji to measure the average pixel intensity values within three distinct regions of interest (ROIs): ROI1: bleached area; ROI2: unbleached area within the cell; and ROI3: background. Normalised FRAP recovery curves and the mobile fractions were calculated using the program easyFRAP (Rapsomaniki et al., 2012).

### Scanning and transmission electron microscopy

For scanning electron microscopy, cells were fixed and processed as already described (Maxeiner et al., 2015; Sechi et al., 2016). Samples were examined with a digital scanning electron microscope (ESEM XL30 FEG; FEI, Hillsboro, OR) using a working distance of 8 mm and an acceleration voltage of 10 kV.

### Statistical analysis

Graphs and statistical tests were done using Prism 8 (GraphPad Software, La Jolla, CA). Differences between sample pairs were analysed using the two-tailed Mann– Whitney nonparametric *U* test. The null hypotheses (the two samples have the same median values, that is, they are not different) were rejected when *p* > 0.5. For the box-and-whiskers plots, the line in the middle of the box indicates the median, the top of the box indicates the 75th quartile, and the bottom of the box indicates the 25th quartile. Whiskers represent the 10th (lower) and 90th (upper) percentiles.

## Results

### LSP1 is essential for efficient migration of J774 mouse macrophages

Since LSP1 directly interacts with F-actin (Wong et al., 2003) and regulates the dynamics of the actin cytoskeleton (Maxeiner et al., 2015), we hypothesised that LSP1 could control cell migration. To this end, we focused on macrophages because LSP1 is essential for another actin-dependent process in this cell type, namely Fcγ receptor-mediated phagocytosis (Maxeiner et al., 2015). After seeding control cells or cells in which LSP1 was down regulated by shRNA (Maxeiner et al., 2015), we imaged cell migration by phase contrast microscopy over a period of 24 hours. Typically, control cells were characterised by a polarised morphology and the formation of large lamellipodia, which developed in the direction of movement (Fig. 1A and Fig. 1SUP). Furthermore, control J774 cells usually travelled large distances (Fig. 1A, C). Conversely, LSP1-deficient J774 cells rarely formed lamellipodia and moved over short distances (Fig. 1B, D and Fig. 1SUP). Consistent with these observations, the average speed of LSP1-deficent J774 cells was significantly smaller than that of control cells (0.01083 µm/sec for LSP1-deficient cells (n=138) vs. 0.02889 µm/sec for control cells (Fig. 1E; n=149). These findings clearly show that LSP1 is essential for efficient migration of J774 macrophages.

**Figure 1.**
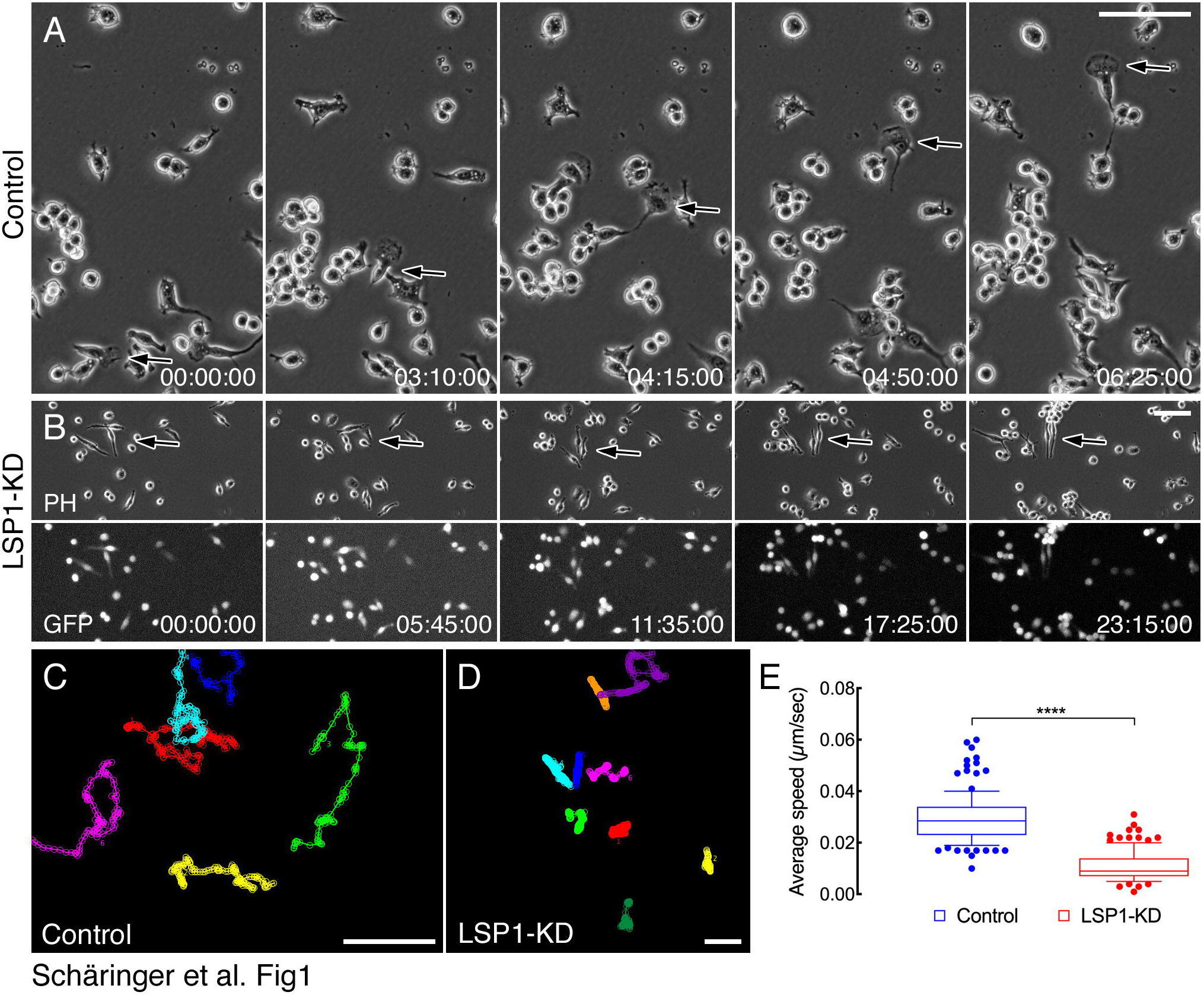
LSP1 is essential for macrophage cell migration. (A, B) Control and LSP1-deficient J774 were cultured on glass coverslips for 24 hours before being imaged by phase contrast microscopy. Control J774 cells characteristically developed large lamellipodia and travelled large distances (arrows in A). By contrast, LSP1-deficient J774 cells (LSP1-KD) did not form lamellipodia and displayed reduced migration (arrows in B). Lower panel in B (GFP) shows the expression of LSP1-specific shRNA. Numbers indicate the elapsed time in hours, minutes and seconds. (C, D) Representative migration tracks of control (C) and LSP1-deficient (D) J774 cells. Scale bars (A-D): 100 µm. (E) Quantification of the average speed of control and LSP1-deficient J774 cells. Box-and-whiskers plots. The line in the middle of the box indicates the median, the top of the box indicates the 75^th^ quartile, and the bottom of the box indicates the 25^th^ quartile. Whiskers represent the 10^th^ (lower) and 90^th^ (upper) percentiles. \*\*\*\**p* < 0.0001.

### LSP1 is essential for normal development of microfilaments, microtubules and focal adhesions

The morphological features and largely decreased migration of LSP1-deficient cells suggest that LSP1 is involved in the organisation of actin and microtubule cytoskeletons as well as cell-substrate adhesion (i.e., focal adhesions). We verified this hypothesis by labelling control and LSP1-deficient J774 cells with anti-tubulin and anti-EB1 antibodies (for assessing microtubule organisation) or fluorescent phalloidin and anti-vinculin antibodies (for assessing microfilaments and focal adhesions, respectively). Using TIRF microscopy, we found that control cells developed a prominent microtubule network characterised by long microtubules emanating from a perinuclear area and projecting toward the cell periphery (Fig. 2A, arrows in inset). As expected, peripheral microtubule ends were labelled with EB1, a plus-end protein that regulates microtubule dynamics (Fig. 2A, arrows in inset). By contrast, the microtubule network in LSP1-deficient cells, which were round and smaller than control cells, was formed by short microtubules (Fig. 2B, arrows in inset). In these cells, we could not find any gross alteration of EB1 distribution (Fig. 2B, arrows in inset). Next, we analysed control and LSP1-deficient J774 cells labelled with Alexa 594-phalloidin and anti-vinculin antibody to visualise actin cytoskeleton and focal adhesion by TIRF microscopy, respectively. Control cells were characterised by a spread and elongated morphology with one or multiple large actin-rich lamellipodia at their periphery (Fig. 2C, green arrowheads). These cells interacted with the substratum via several elongated focal adhesions (Fig. 2C, arrows). At variance with these morphological features, LSP1-deficent cells were smaller and round with no or a single small actin-rich lamellipodium (Fig. 2D, red arrowhead). Focal adhesions in these cells were strongly reduced in size and number and showed a rounded shape (Fig. 2D, arrows). Overall, these findings demonstrate that LSP1 is essential for the normal development of microfilaments, microtubules and focal adhesions. They also suggest that LSP1-dependent regulation of cell migration is exerted via the control of these cytoskeletal structures.

**Figure 2.**
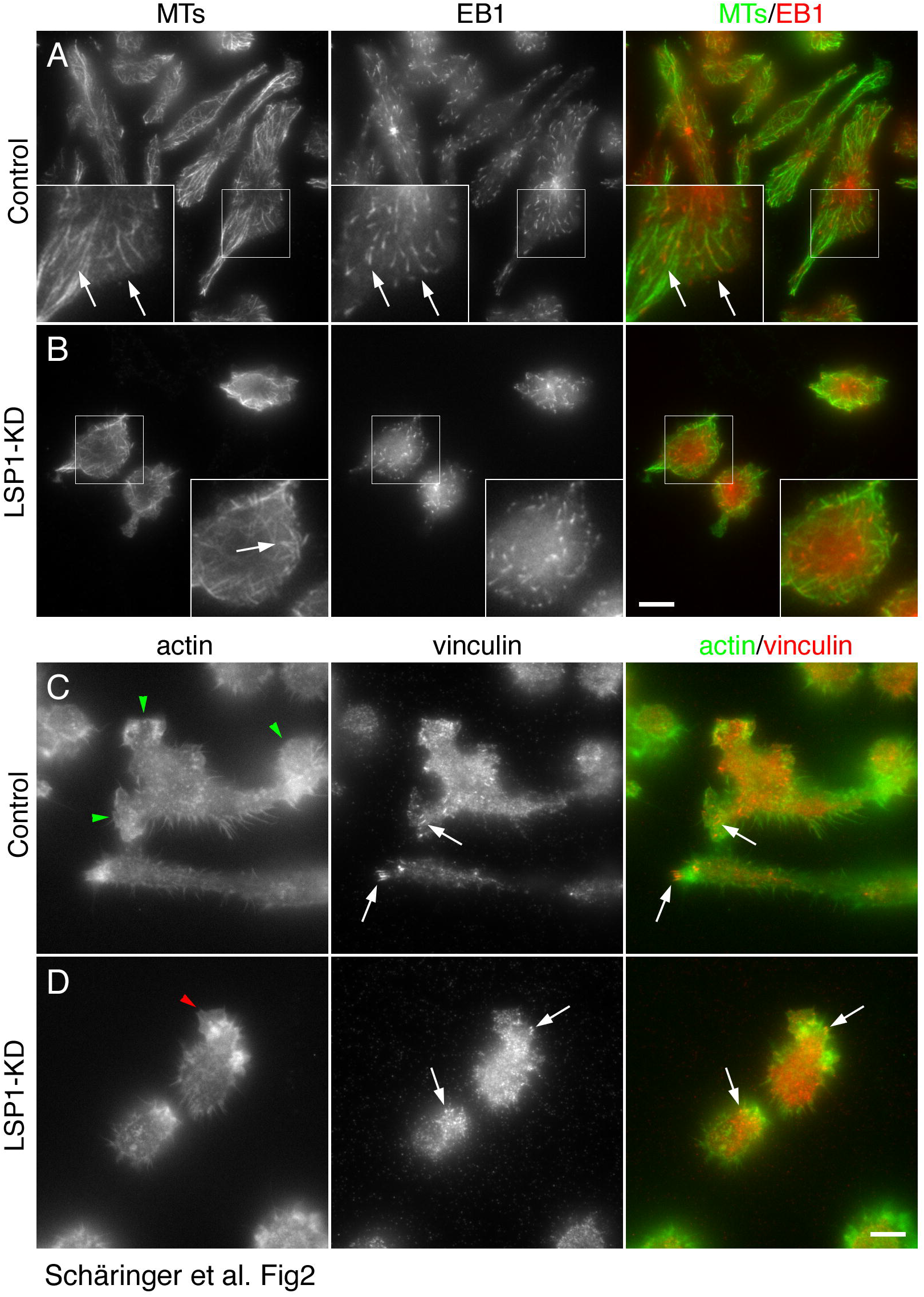
LSP1 is essential for normal development of microfilaments, microtubules and focal adhesions. (A, B) Microtubule and EB1 cytoplasmic distribution in control and LSP1-deficient (LSP1-KD) J774 cells. Following fixation, microtubules and EB1 were visualised using anti-tubulin and anti-EB1 antibodies, respectively. Samples were analysed by TIRF microscopy. In control cells (A), microtubules were well-developed typically originating from a perinuclear area and projecting towards cell periphery (left panel, arrows in inset). EB1 characteristically localised at the peripheral tips of microtubules (central and right panels, arrows in insets). LSP1-deficient cells acquired a rounded shape and were characterised by shorter microtubules (B, left panel, arrow in inset). EB1 localisation was not grossly changed. Right panels show merged microtubule (shown in green) and EB1 (shown in red) images. Boxes indicate the areas enlarged in the insets. Scale bar: 10 µm. (C, D) Actin cytoskeleton and focal adhesion distribution in control and LSP1-deficient (LSP1-KD) J774 cells. The actin cytoskeleton was visualised using fluorescent phalloidin, whereas focal adhesions were detected using anti-vinculin antibodies. Samples were analysed by TIRF microscopy. Control cells formed one or more large actin-rich lamellipodia (C, left panel, green arrowheads). Conversely, LSP1-deficient cells formed only small lamellipodia (D, left panel, red arrowhead). Focal adhesions were also altered in LSP1-deficient cells, which formed fewer and smaller focal adhesions than control cells (arrows in central and right panels). Right panels show merged actin (shown in green) and vinculin (shown in red) images. Scale bar: 10 µm.

### LSP1 is essential for the regulation of focal adhesion dynamics

Focal adhesions are highly dynamic structures whose spatial and temporal regulation is essential for cell migration (Sechi and Wehland, 2004; Zamir and Geiger, 2001). The impaired cell migration and formation of focal adhesions in LSP1-deficent J774 cells suggests that LSP1 plays an important role in the control of focal adhesion dynamics. To test this assumption, we engineered control and LSP1-deficient J774 cells to express RFP-tagged zyxin, a focal adhesion component, to visualise focal adhesion dynamics using TIRF microscopy (Gamper et al., 2016; Sechi et al., 2016) and analysed their dynamics using a dedicated algorithm (Würflinger et al., 2011). The initial examination of time-lapse sequences revealed that focal adhesions in control cells were highly dynamic assembling or disassembling within short time periods (arrows in Fig. 3A and corresponding video). On the contrary, focal adhesions in LSP1-deficient cells appeared to be less dynamic requiring longer time periods to assemble and disassemble (arrows in Fig. 3B and corresponding video). The quantification of several focal adhesion parameters confirmed the impression provided by visually inspecting time-lapse sequences. Precisely, for both growing or shrinking focal adhesions, the change of the area over time, a proxy for assembling and disassembly rates, was faster for focal adhesions in control cells than in LSP1-deficient cells (Fig. 3C, D). Accordingly, the assembly and disassembly rates of focal adhesions in LSP1-deficient cells were significantly lower than the corresponding parameters for control focal adhesions (Fig. 3E). Furthermore, the average area was significantly reduced in focal adhesions in LSP1-deficient cells (Fig. 3F), whereas we could not see any difference in their shape (elongation index, Fig. 3G). Finally, the focal adhesion movement relative to the substratum (focal adhesion speed) was also significantly impaired in LSP1-deficient cells (Fig. 3H). These findings clearly show that LSP1 regulates cell migration via the modulation of focal adhesion formation and dynamics.

**Figure 3.**
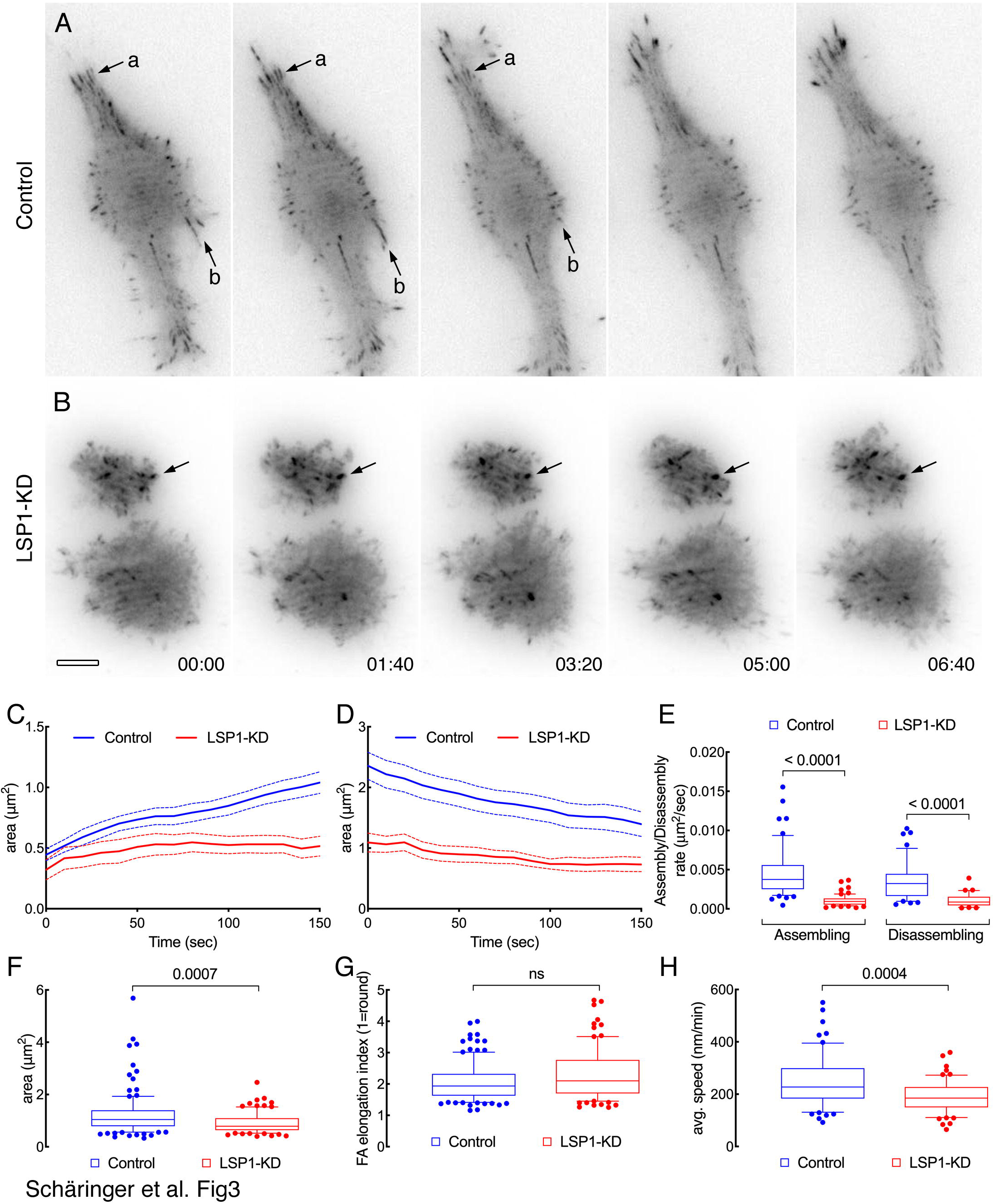
LSP1 is essential for the regulation of focal adhesion dynamics. (A, B) Representative time-lapse images showing focal adhesion dynamics in control (A) and LSP1-deficient (B; LSP1-KD) J774 cells. Focal adhesions were visualised using RFP-zyxin and images were acquired by TIRF microscopy. Note the faster turnover of focal adhesions in control cells (arrows in A) compared to focal adhesions in LSP1-deficient cells (arrows in B). Numbers indicate the elapsed time in minutes and seconds. Scale bar: 10 µm. (C-H) Quantification of focal adhesion parameters. In the box-and-whiskers plots the line in the middle of the box indicates the median, the top of the box indicates the 75^th^ quartile, and the bottom of the box indicates the 25^th^ quartile. Whiskers represent the 10^th^ (lower) and 90^th^ (upper) percentiles. ns: non-significant.

### LSP1-myosin1e binary complex is essential for the regulation of cell migration

We have demonstrated that LSP1 directly interacts with myosin 1e through a non-canonical SH3-binding site. Moreover, downregulating LSP1 or blocking its interaction with myosin1e results in severely impaired Fcγ receptor-mediated phagocytosis and the inhibition of actin accumulation and lamellipodia formation around the particles to be internalised (Maxeiner et al., 2015). Since LSP1 deficiency impairs cell migration, we reasoned that the LSP1-myosin1e binary complex could play an important role in the regulation of J774 cell migration. To experimentally verify this hypothesis, we scrutinised the migration of J774 cells expressing the deletion mutant LSP1-ΔSBS, which cannot bind to myosin1e (Maxeiner et al., 2015), by phase contrast microscopy for 24 hours. As control, we used J774 cells re-expressing full-length LSP1 (rescue). As expected, cells in which LSP1 was re-expressed did not shown any sign of migration defect highly resembling control J774 cells (Fig. 4B, C, E). By contrast, it was immediately evident that cells expressing LSP1-ΔSBS travelled very short distances (Fig. 4A, D) and moved at a significantly reduced speed (Fig. 4E). Remarkably, the motile phenotype and the speed of cells expressing LSP1-ΔSBS was undistinguishable from LSP1-deficient cells (compare with Fig. 1). These observations clearly demonstrate that the binary complex between LSP1 and myosin1e is essential for efficient J774 migration.

**Figure 4.**
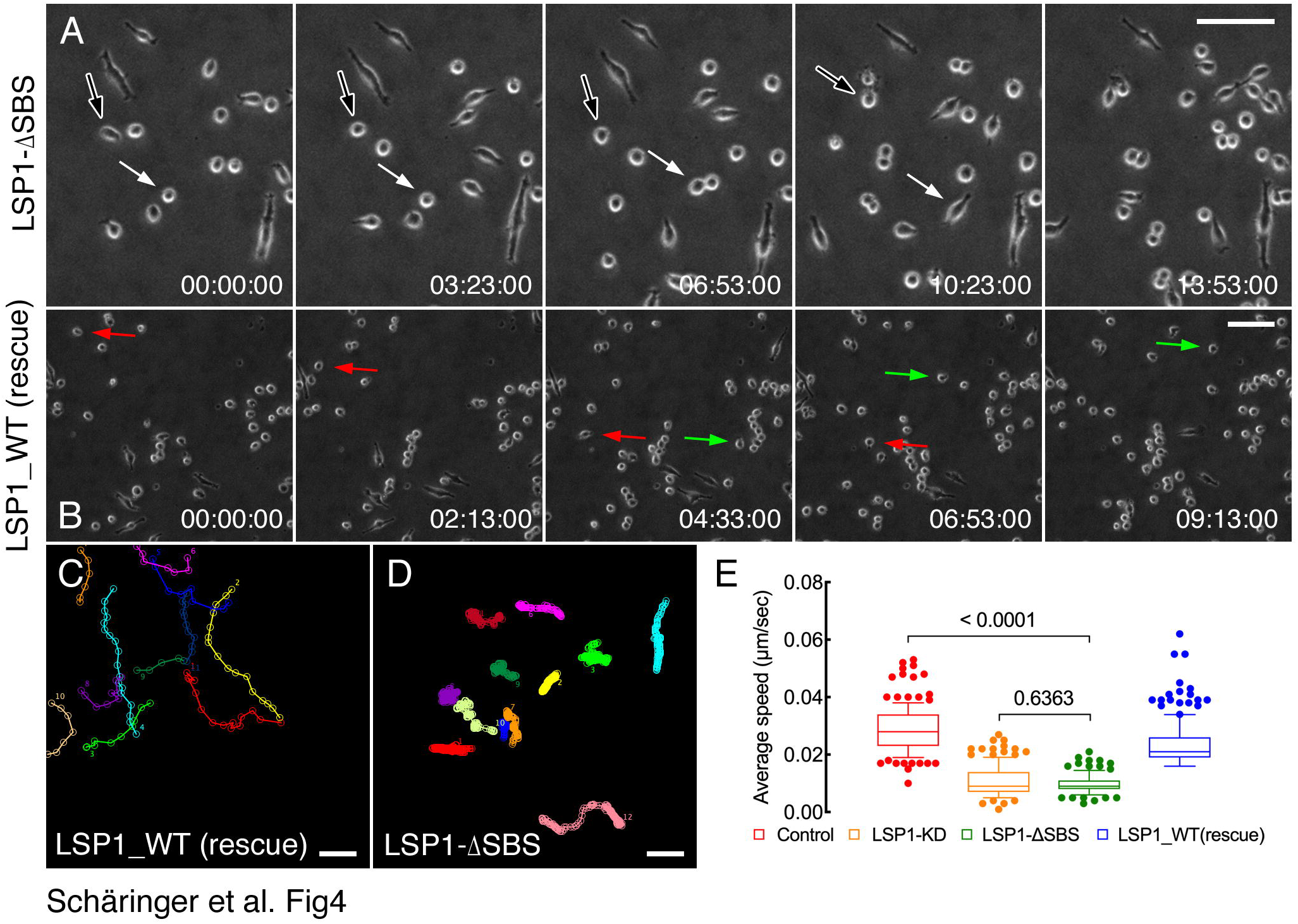
LSP1-myosin1e interaction is essential for J774 cell migration. (A, B) J774 cells expressing the LSP1-ΔSBS (A) or full-length LSP1 (B; LSP1_WT rescue) were cultured on glass coverslips for 24 hours before being imaged by phase contrast microscopy. J774 cells in which the expression of full-length LSP1 was restored (rescued; arrows in B) moved at faster speed compared to cells expressing the LSP1 mutant LSP1-ΔSBS, which is unable to interact with myosin 1e (arrows in A). Numbers indicate the elapsed time in hours, minutes and seconds. (C, D) Representative migration tracks of rescue (C) and LSP1-ΔSBS (D) J774 cells. Scale bars (A-D): 100 µm. (E) Quantification of the average speed of LSP1-ΔSBS and rescue J774 cells. For easier comparison, the average speeds of control and LSP1-deficient cells (data from Fig. 1) are also plotted. Box-and-whiskers plots. The line in the middle of the box indicates the median, the top of the box indicates the 75^th^ quartile, and the bottom of the box indicates the 25^th^ quartile. Whiskers represent the 10^th^ (lower) and 90^th^ (upper) percentiles.

### LSP1-myosin 1e binary complex is necessary for lamellipodia activity

One of the earliest events of cell migration is the formation and stabilisation of one lamellipodium in the direction of movement. Because cell migration is severely impaired in LSP1-deficient cells and in cells in which LSP1 cannot interact with myosin1e, we decided to determine whether lamellipodia activity is compromised in these cell types. The closer inspection of time-lapse sequences at high magnification revealed that motile control cells frequently form one large and persistent lamellipodium in the direction of movement (Fig. 5A and corresponding video). LSP1-deficient cells and cells expressing LSP1-ΔSBS greatly differed from this phenotype in that they maintained a round morphology and never formed a large and persistent lamellipodium (Fig. 5B, C). These cells were rather characterised by the formation of small lamellipodia that formed around their periphery (Fig. 5B, C and corresponding video). Consistent with this visual examination, we found that the frequency of lamellipodia formation and its width were significantly lower in LSP1-deficient cells and cells expressing LSP1-ΔSBS (Fig. 5D, F). Interestingly, the speed of lamellipodia spreading was significantly reduced in LSP1-ΔSBS cells, but not in LSP1-deficient cells (Fig. 5E) possibly due to a large data variability. Collectively, these findings demonstrate that LSP1 and myosin1e are indispensable for efficient lamellipodia activity.

**Figure 5.**
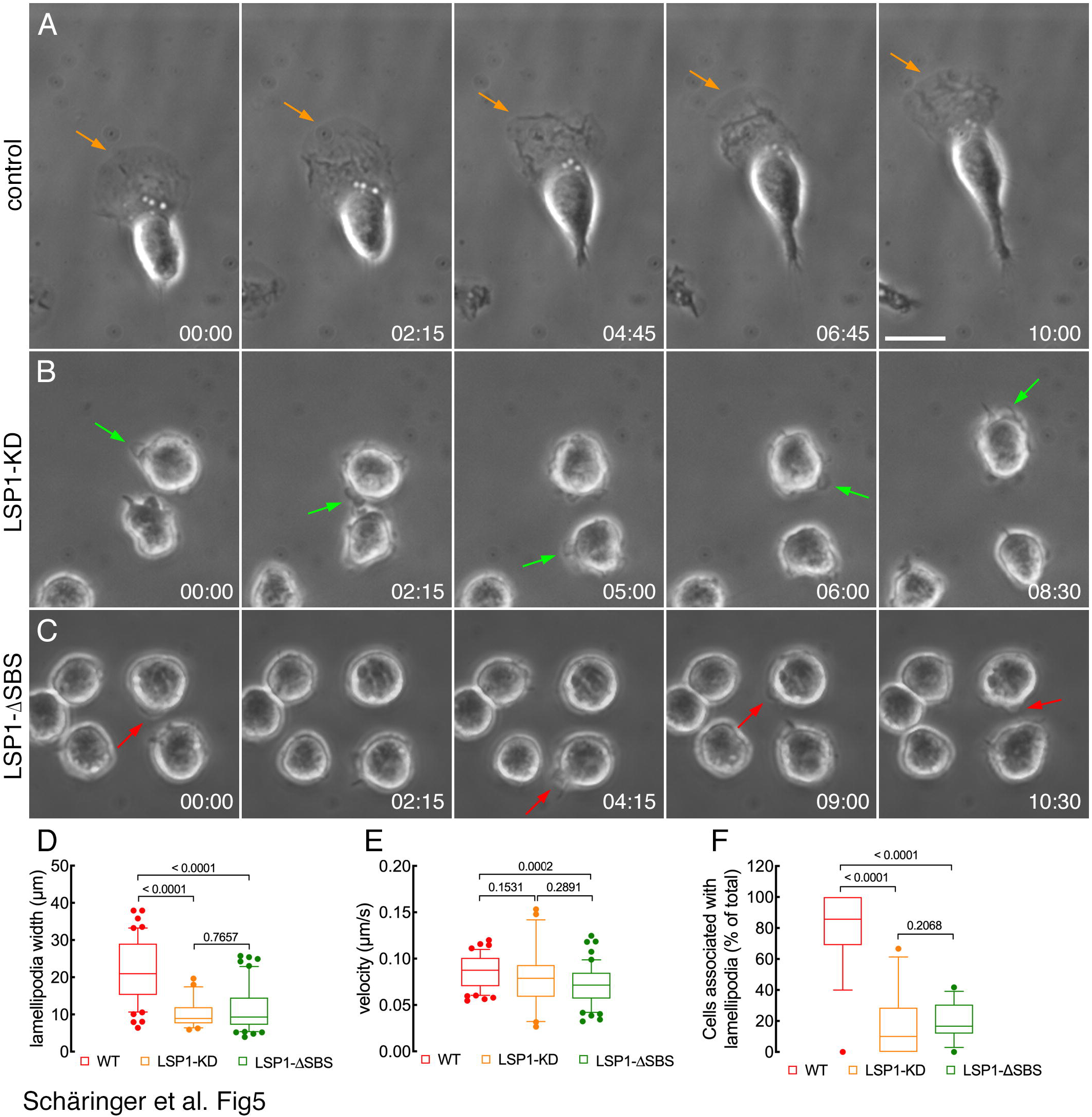
LSP1-myosin1e complex is essential for lamellipodia formation and dynamics. (A-C) Time-lapse images showing lamellipodia morphology and dynamics in control (A), LSP1-deficient (B; LSP1-KD) and J774 cells expressing the LSP1 deletion mutant LSP1-ΔSBS (C). Polarised control cells typically formed one large and very dynamic lamellipodium in the direction of movement (arrows in A). By contrast, LSP1-deficient cells or cells expressing an LSP1 mutant unable to interact with myosin 1e acquired a rounded morphology and formed very small lamellipodia around their periphery (arrows in B, C). Numbers indicate the elapsed time in minutes and seconds. Scale bar: 10 µm. (D-F) Box-and-whiskers plots showing the quantification of width, spreading velocity and frequency of lamellipodia formation. The line in the middle of the box indicates the median, the top of the box indicates the 75^th^ quartile, and the bottom of the box indicates the 25^th^ quartile. Whiskers represent the 10^th^ (lower) and 90^th^ (upper) percentiles.

### LSP1-myosin1e interaction regulates the dynamics and kinetics of LSP1

Next, we determined whether the interaction with myosin1e affected the dynamics and kinetics of LSP1. We initially visualised LSP1 localisation and dynamics by TIRF microscopy over a period of 10-15 minutes. In control cells and cells re-expressing full-length LSP1 (rescue), LSP1 was highly dynamic often localising to lamellipodia (asterisk in Fig. 6A and C; see also corresponding video) and to filamentous-like and focal adhesions-like structures (arrows in Fig. 6A and C). In cells expressing LSP1-ΔSBS, LSP1 appeared to be less dynamics and was concentrated at the perinuclear area and at very small lamellipodia (arrows in Fig. 6B; see also corresponding video). To corroborate the visual impression that LSP1 was less dynamic when unable to interact with myosin1e, we determined its kinetics using FRAP microscopy. As shown in Fig. 6D and E, both the time-course of the fluorescence recovery and the mobile fraction of LSP1 in control cells and cells re-expressing full-length LSP1 (rescue) were indistinguishable. Conversely, the time-course of the fluorescence recovery and the mobile fraction of LSP1-ΔSBS was significantly reduced (Fig. 6, D, E). Thus, our findings suggest that efficient LSP1 dynamics and kinetics depend on its interaction with myosin1e.

**Figure 6.**
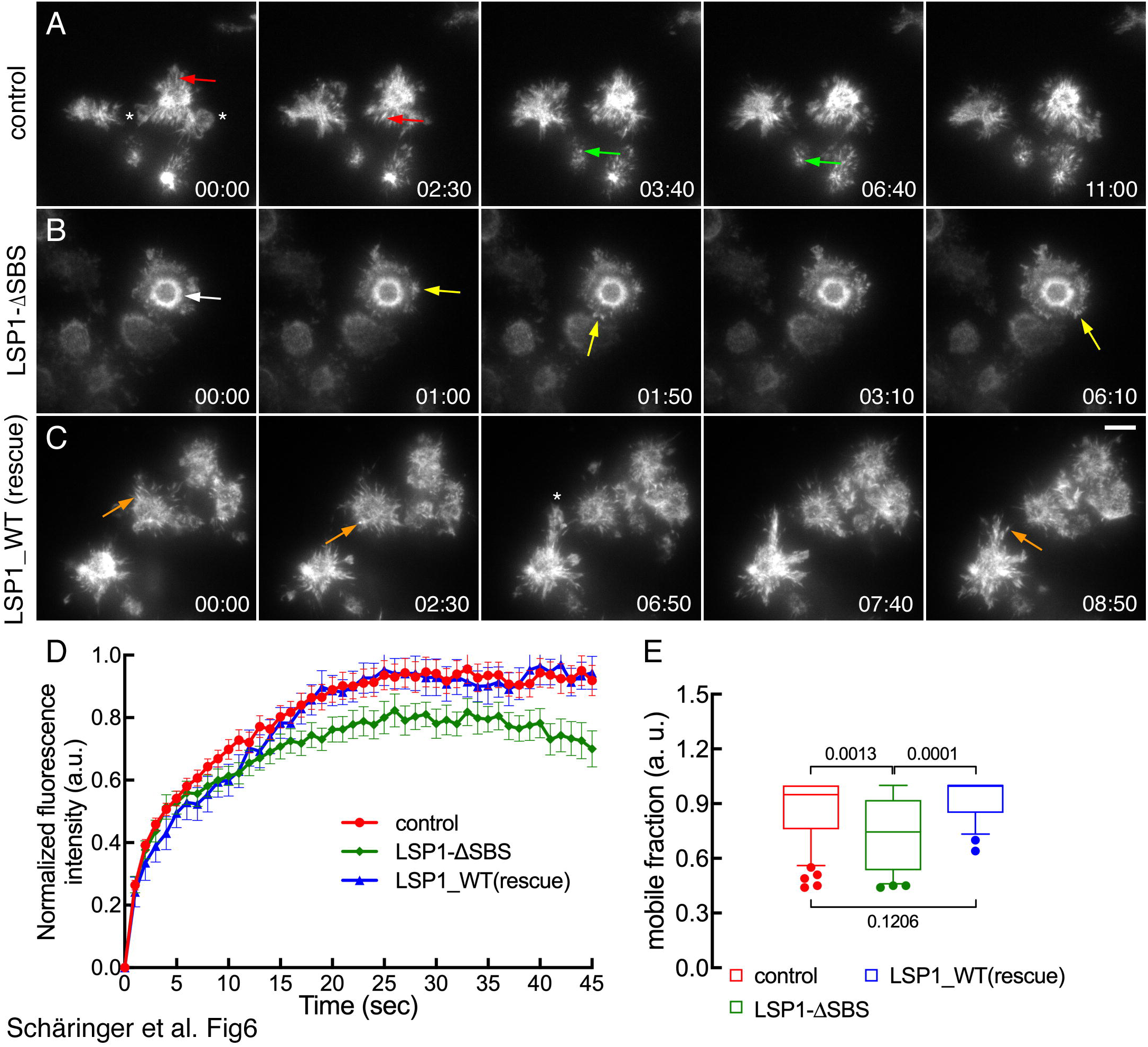
LSP1-myosin1e interaction regulates the dynamics and kinetics of LSP1. (A-C) TIRF imaging showing the dynamics of LSP1 in control J774 cells (A), cells expressing the LSP1 mutant unable to interact with myosin 1e (LSP1-ΔSBS; B) or full-length LSP1 (rescue; C). Note the intense dynamics of LSP1 in control (A) and rescue cells (C) and the localisation of LSP1 to filamentous-like structures (red arrows in A) and focal adhesions-like structures (green arrows in A, orange arrows in C). Stars indicate lamellipodia. In cells expressing LSP1-ΔSBS, LSP1 robustly localises around the nucleus (white arrow in B) and at very small lamellipodia (yellow arrows in B). Numbers indicate the elapsed time in minutes and seconds. Scale bar: 10 µm. (D, E) Quantification of the fluorescence recovery (D) and mobile fraction (E) of LSP1. Error bars show s.e.m. In the box-and-whiskers plots, the line in the middle of the box indicates the median, the top of the box indicates the 75^th^ quartile, and the bottom of the box indicates the 25^th^ quartile. Whiskers represent the 10^th^ (lower) and 90^th^ (upper) percentiles.

### LSP1-myosin1e binary complex is essential for the regulation of focal adhesion dynamics

The above findings provide clear evidence that LSP1 deficiency severely impairs cell migration, focal adhesion dynamics and lamellipodia activity. Moreover, the interaction of LSP1 with myosin1e is essential for the regulation of cell migration and lamellipodia formation. Based on these observations and given the importance of focal adhesion dynamics for cell migration, we sought to determine whether LSP1-myosin1e binary complex also plays a role in the regulation of focal adhesion dynamics. For this purpose, we genetically modified J774 expressing LSP1-ΔSBS or full-length LSP1 (rescue) with turquoise-zyxin to visualised focal adhesion dynamics using TIRF microscopy.

Similar to LSP1-deficient cells, LSP1-ΔSBS cells were characterised by few and small focal adhesions with a slower turnover (Fig. 7B and corresponding video). As expected, focal adhesions in cells re-expressing full-length LSP1 were similar to those in control cells (Fig. 7A and corresponding video; compare with Fig. 3A). Detailed analysis of focal adhesion dynamics using a dedicated algorithm supported this visual impression showing that assembly and disassembly rates as well as size were significantly reduced in LSP1-ΔSBS cells (Fig. 7C-H). Consistent with these observations, we found a reduced amount of vinculin in the actin cytoskeleton fraction of LSP1-deficient and LSP1-ΔSBS cells (Fig. 8).

**Figure 7.**
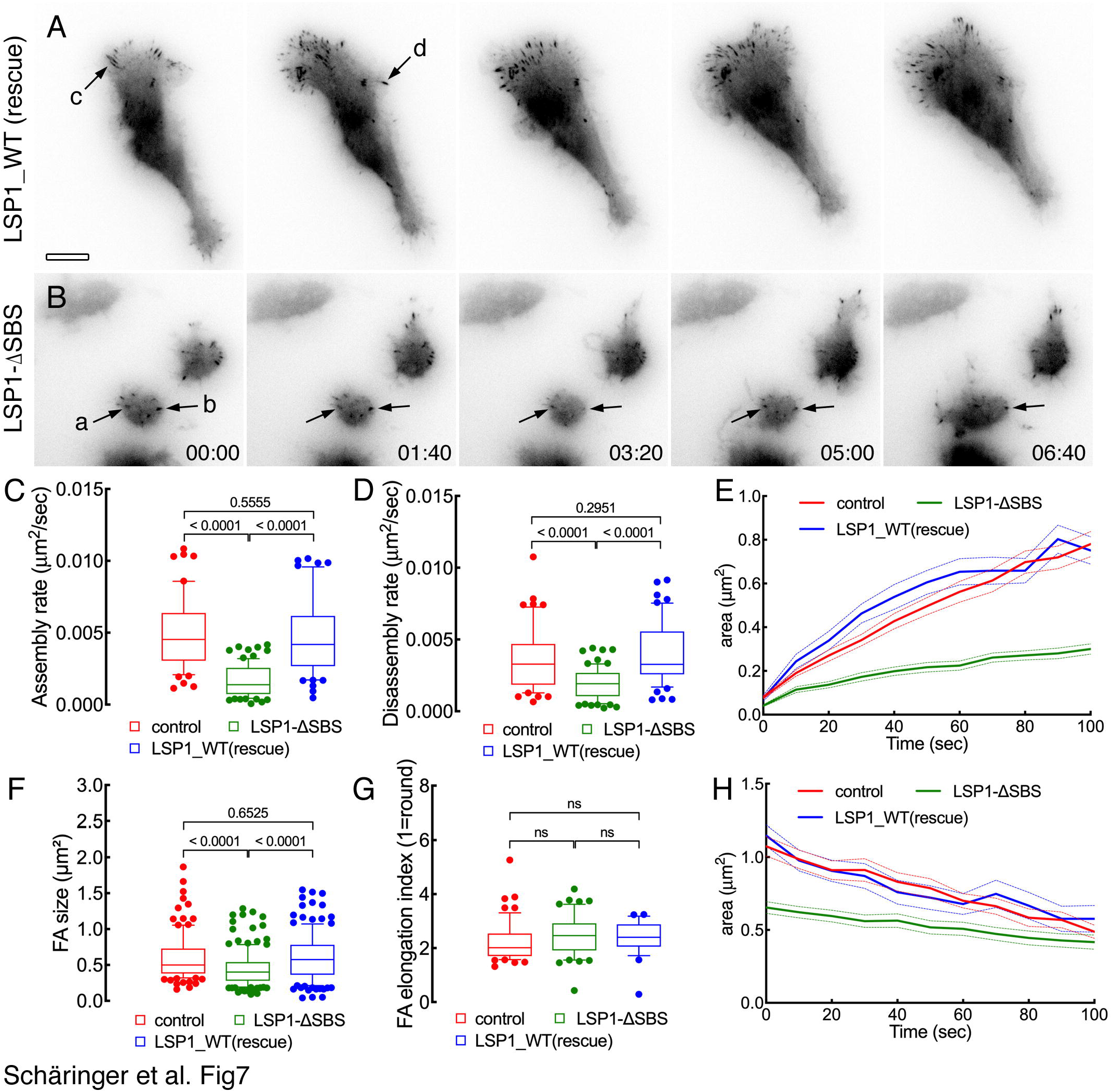
LSP1-myosin1e interaction is essential for efficient focal adhesion dynamics in J774 cells. (A, B) J774 cells expressing full-length LSP1 (A; rescue) or the LSP1 mutant unable to interact with myosin 1e (B; LSP1-ΔSBS) were transfected with RFP-zyxin to visualise focal adhesions and images were acquired by TIRF microscopy. Note the faster turnover of focal adhesions in cells expressing full-length LSP1 (arrows in A) compared to focal adhesions in cells expressing the LSP1 mutant unable to interact with myosin 1e (arrows in B). Numbers indicate the elapsed time in minutes and seconds. Scale bar: 10 µm. (C-H) Quantification of focal adhesion parameters. In the box-and-whiskers plots the line in the middle of the box indicates the median, the top of the box indicates the 75^th^ quartile, and the bottom of the box indicates the 25^th^ quartile. Whiskers represent the 10^th^ (lower) and 90^th^ (upper) percentiles. ns: non-significant.

**Figure 8.**
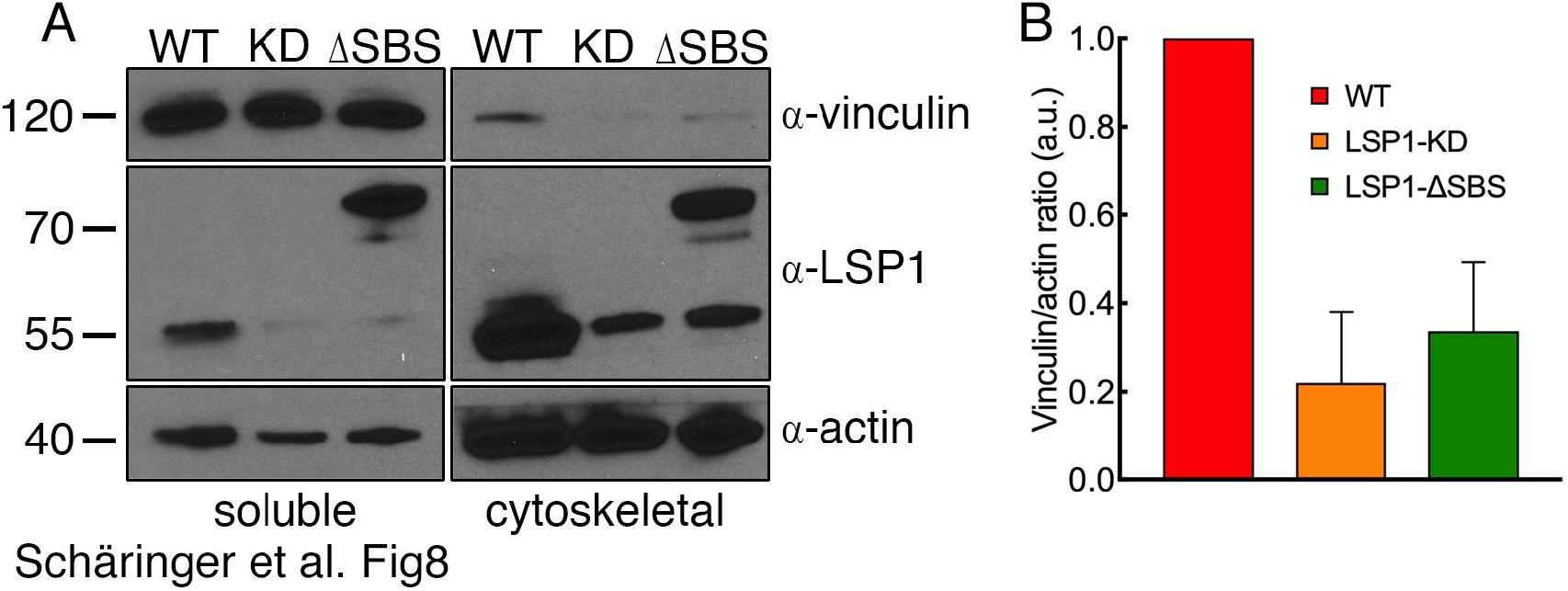
LSP1-myosin1e interaction regulates the amount of actin-associated vinculin. (A) Soluble and cytoskeletal fractions from control (WT), LSP1-deficient (LSP1-KD) and J774 cells expressing the LSP1-ΔSBS mutant (unable to interact with myosin1e) resolved by SDS-PAGE and probed with antibodies against vinculin, LSP1 and actin, which served as loading control. Numbers on the left side indicate MW markers in kDa. Note the reduced amount of vinculin recovered in the cytoskeleton fraction of LSP1-deficient cells and cells expressing the deletion mutant LSP1-ΔSBS. (B) Quantification of vinculin/actin ratio in cytoskeleton fraction. Vinculin/actin ratio for control cells (WT) was set to 1. Columns show mean ± s.e.m (n=3).

Next, as a complementary approach to demonstrate the role of LSP1-myosin1e binary complex in the regulation of focal adhesions, we analysed zyxin kinetics by fluorescence recovery after photobleaching. To this end, focal adhesions in control, LSP1-deficient, LSP1-ΔSBS and cells re-expressing full-length LSP1 (rescue) were bleached and the recovery of zyxin fluorescence within the bleached areas was monitored over a period of several minutes. According to the above analysis of focal adhesion dynamics, zyxin kinetics was not significantly different in control cells and cells re-expressing full-length LSP1 (Fig. 9A, D-F). Conversely, zyxin recovery was much slower and less complete at focal adhesions in LSP1-deficient and LSP1-ΔSBS cells (Fig. 9B, C, E, F). Taken together, these findings demonstrate that the LSP1-myosin1e binary complex is required for the regulation of focal adhesion dynamics.

**Figure 9.**
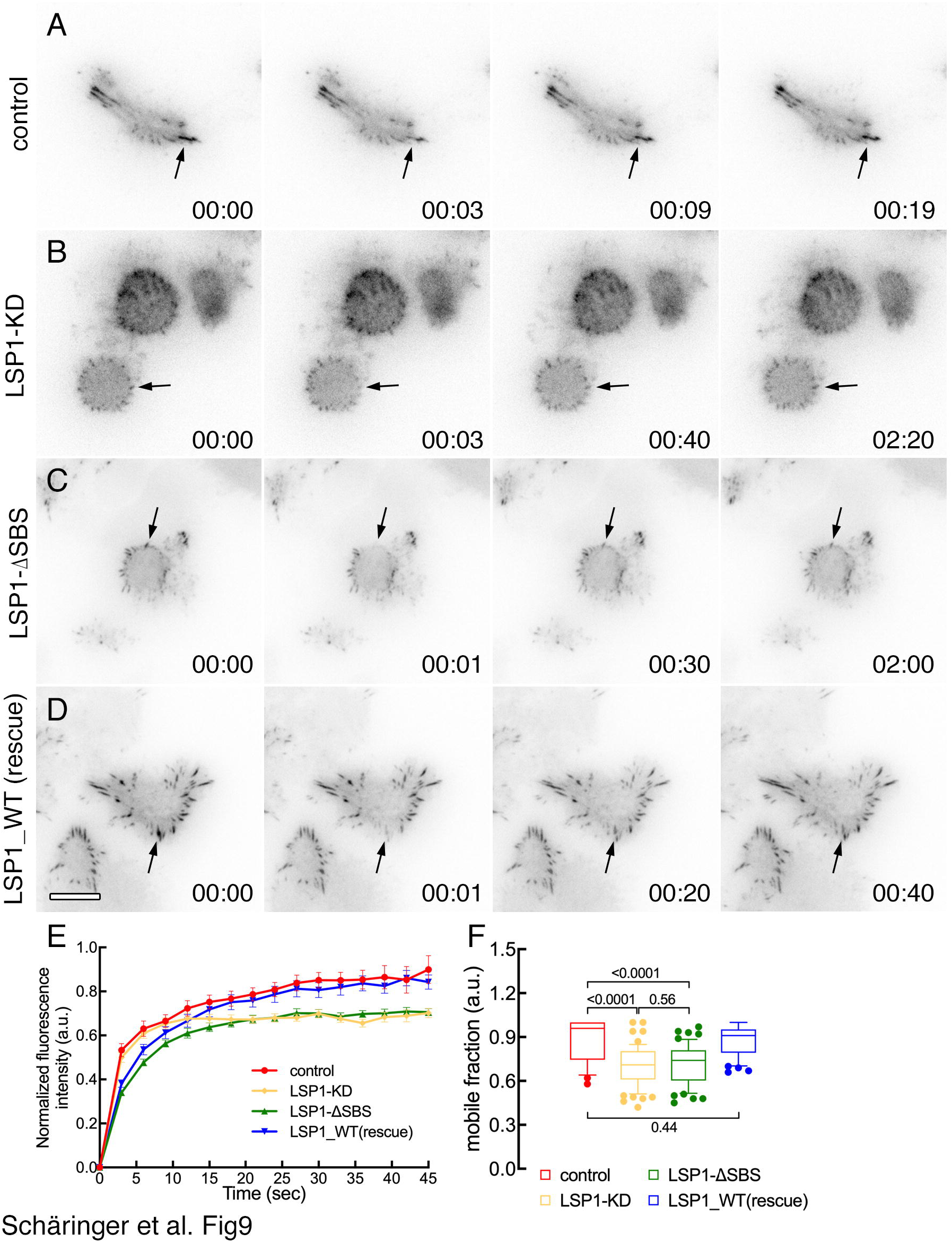
LSP1-myosin1e interaction is essential for efficient kinetics of zyxin at focal adhesions. (A-D) Control J774 cells (A), LSP1-deficient cells (LSP1-KD; B), cells expressing the LSP1 mutant unable to interact with myosin 1e (LSP1-ΔSBS; C) or full-length LSP1 (rescue; D) were stably transfected with RFP-zyxin to visualise focal adhesions. RFP-zyxin kinetics was determined by fluorescence recovery after photobleaching. Arrows in A-D point to bleached focal adhesions. Note the slower recovery of the fluorescence signal at bleached focal adhesions in LSP1-deficient cells (B) and in cells expressing the LSP1 mutant unable to interact with myosin1e (C). (E, F) Quantification of the fluorescence recovery (E) and mobile fraction (F) of RFP-zyxin. Error bars show s.e.m. In the box-and-whiskers plots, the line in the middle of the box indicates the median, the top of the box indicates the 75^th^ quartile, and the bottom of the box indicates the 25^th^ quartile. Whiskers represent the 10^th^ (lower) and 90^th^ (upper) percentiles.

## Discussion

Deciphering the mechanisms of cell migration is essential for fully understanding this process in normal and pathological conditions. In this study, we demonstrate that LSP1 is essential for efficient cell migration in J774 macrophages. LSP1 controls this process through the regulation of the formation and dynamics of lamellipodia and focal adhesions. In this context, the interaction of LSP1 with the molecular motor myosin1e is indispensable for LSP1 function in cell migration since inhibiting the formation of LSP1-myosin1e binary complex severely impairs cell migration, formation and dynamics of lamellipodia and focal adhesions. These findings provide novel insights into the regulation of cell migration in immune cells and demonstrate a pivotal role for the LSP1-myosin1e binary complex in the regulation of this process.

Previous investigations have addressed the role of LSP1 in cell migration in a variety of cell types. For instance, LSP1 deficiency significantly reduces the migration of dendritic cells induced by gp120 (Anand et al., 2009; Prasad et al., 2012). Likewise, LSP1^−/-^ neutrophils show both impaired migration and chemotactic response (Hannigan et al., 2001). Using a complementary approach, it was shown that the expression of LSP1 to physiological levels in a myeloid cell line, which does not express LSP1, results in a significant enhancement of cell migration (Li et al., 2000). Notably, large lamellipodia are rarely observed in LSP1^−/-^ cells, which rather form small and transient lamellipodia (Hannigan et al., 2001). In this study, we have deepened our knowledge of LSP1 function demonstrating that its deficiency severely impairs the formation of lamellipodia, focal adhesion dynamics and cell migration in macrophages. Hence, LSP1 unequivocally plays a key role in the regulation of cell migration in different cellular systems.

For the sake of clarity, it should be mentioned that other studies show that the loss, or reduced expression, of LSP1 increases rather than reduces cell migration. However, this effect has been described in pathological situations such as in T-cells derived from patients with rheumatoid arthritis and in hepatoma cells (Hwang et al., 2015; Koral et al., 2015). Since, in hepatoma cells, a classical scratch assay and not single-cell tracking was used to study cell migration, it is not possible to rule out that the described increase of cell migration was not due to the increase of cell proliferation observed in LSP1^−/-^ cells. In addition, the marginal (1.36-fold) increase of migration of LSP1^−/-^ neutrophils could be observed when cells were seeded only on fibrinogen, but not on other substrates (Wang et al., 2002). Hence, when interpreting these studies ascribing a negative regulatory effect on cell migration to LSP1, experimental parameters that may impact on this effect must be considered in order to avoid inaccurate interpretations.

Although it is known that LSP1 interacts with actin filaments (Klein et al., 1990; Wong et al., 2003; Zhang et al., 2001), the mechanisms underlying LSP1 function in the regulation of actin-driven processes are, as yet, poorly defined. Before discussing potential scenarios describing LSP1 function, our previous study (Maxeiner et al., 2015) and the findings described here clearly indicate that one possible mechanism for LSP1 function requires its interaction with myosin1e. In this context, we envisage that LSP1 and myosin1e will synergise to activate signalling pathways specific for actin cytoskeleton remodelling. How does, then, this binary complex regulate cell migration and focal adhesion dynamics? It has been shown that LSP1 is a component of a complex including WASP and the Arp2/3 complex (Prasad et al., 2012). LSP1 also co-precipitates with regulators of myosin II such as calmodulin and the myosin light chain kinase (Cervero et al., 2018). Since WASP and the Arp2/3 complex are essential for actin filament nucleation in lamellipodia and myosin II activity controls actin cytoskeleton contraction and cell body displacement required for cell migration (Blanchoin et al., 2014), it is reasonable to envisage a signalling scenario in which LSP1 controls both actin filament nucleation and cell body translocation (Fig. 10). Our present study supports this hypothesis since in LSP1-deficient cells both lamellipodia formation and cell migration are severely impaired.

**Figure 10.**
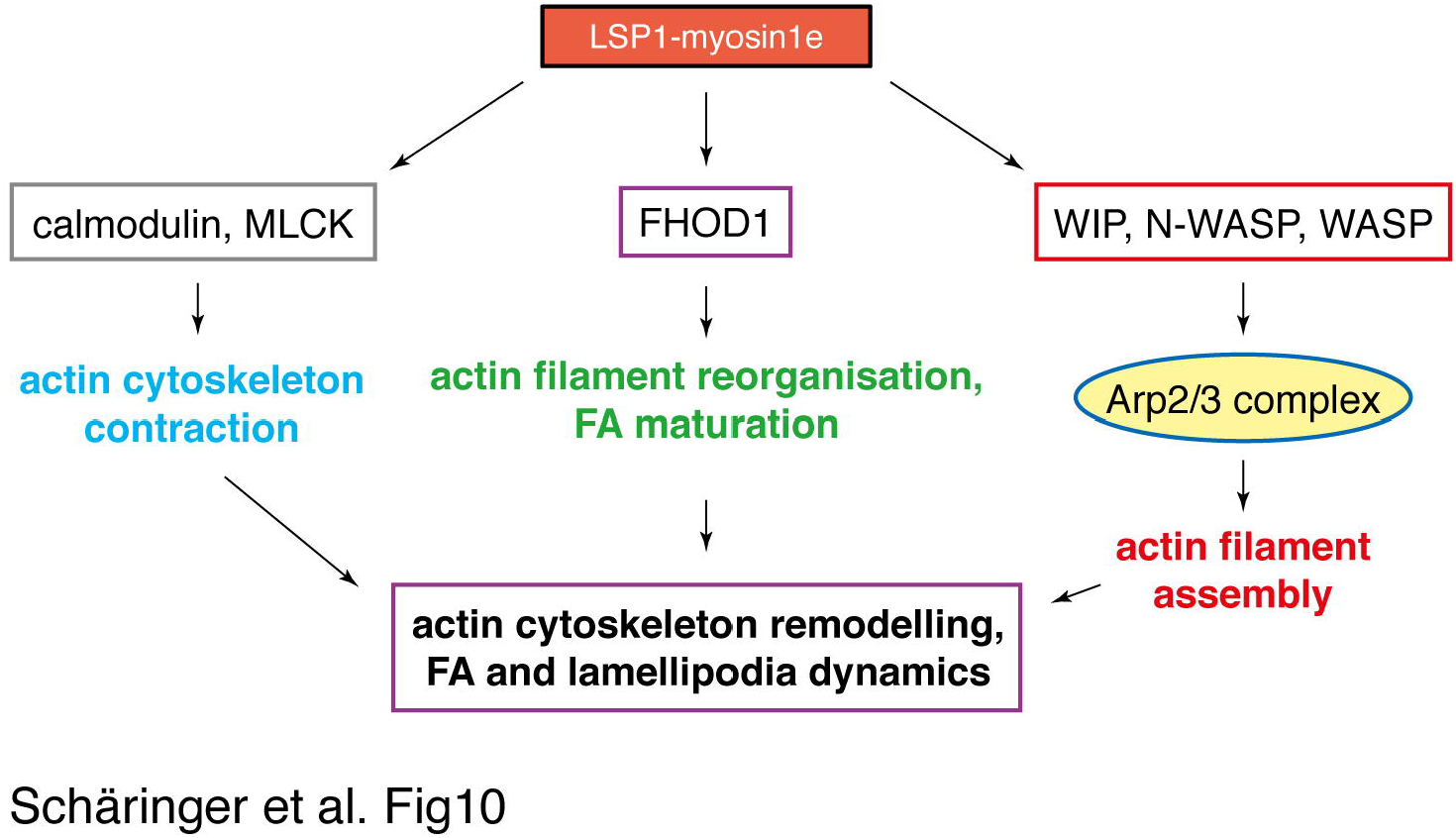
Schematic model for the role of LSP1-myosin1e complex during cell migration and adhesion. The LSP1-myosin1e binary complex could regulate cell adhesion and migration via three different, but not necessarily functionally independent, signalling pathways. The LSP1-myosin1e complex could regulate actin cytoskeleton contraction through calmodulin and myosin light chain kinase, which modulate myosin II activity. Actin filament reorganisation and FA maturation could be controlled via FHOD1. Finally, the LSP1-myosin1e could also impinge on actin filament assembly upon the regulation of WASP family proteins and the Arp2/3 complex. The functional integration of these three signalling pathways will efficiently regulate actin cytoskeleton remodelling, FA and lamellipodia dynamics leading to efficient cell migration.

It is interesting to note that targeting myosin1e to mitochondria induces the accumulation at the surface of these organelles of actin filaments and activators of the Arp2/3 complex WIP and N-WASP (Cheng et al., 2012). Furthermore, in yeast, a type I myosin has been involved in actin filament formation induced by WASP and the Arp2/3 complex (Sirotkin et al., 2005). These studies suggest a role for myosin1e in the formation of actin filaments. This role for myosin1e is further supported by two additional studies showing that lamellipodia formation and stability is impaired in myosin1e-deficient cells (Gupta et al., 2013; Tanimura et al., 2016). It is also important to note that cells expressing a variant of myosin1e lacking the SH3 domain fail to form mature focal adhesions (Gupta et al., 2013). Consistent with these studies, we have demonstrated that lamellipodia and focal adhesion formation are impaired in LSP1-deficent cells and in cells expressing a variant of LSP1 that cannot interact with myosin 1e. Thus, it is conceivable that LSP1 and myosin1e work in concert to regulate actin filament assembly through WASP-family proteins and the Arp2/3 complex (Fig. 10). In this context, it should also be taken into account that myosin 1e has been found in actin-rich structures containing FHOD1 (Gupta et al., 2013). In mammalian cells, FHOD1 stimulates the formation of stress fibres and cell migration and is recruited to integrin clusters (Iskratsch et al., 2013; Koka et al., 2003; Schulze et al., 2014). Moreover, FHOD1 knockdown impairs cell spreading and focal adhesion maturation (Iskratsch et al., 2013). In *Drosophila*, the mutation of the FHOD1 homologue Knittrig results in smaller macrophages, which show reduced migration (Lammel et al., 2014). These observations are, again, in agreement with our findings showing a similar phenotype in LSP1-deficent cells and in cells expressing a variant of LSP1 that cannot interact with myosin1e. Hence, we propose that the LSP1-myosin1e binary complex exert its function also via FHOD1 (Fig. 10).

Finally, it is important to note that the prominence of LSP1 in the regulation of cell migration is further supported by studies on pathogen-cell interactions. It has been clearly demonstrated that several types of pathogens such as *Listeria monocytogenes* and vaccinia viruses have developed elegant strategies to subvert fundamental steps of actin cytoskeleton remodelling for their spreading and survival (Geese et al., 2002; May et al., 1999; Pust et al., 2005; Way, 1998). In this context, it has been shown that the human immunodeficiency virus can induce migration of dendritic cells (Anand et al., 2009). These viruses control dendritic cell migration by the binding of their envelope protein gp120 to DC-SIGN on dendritic cells (Prasad et al., 2017; Prasad et al., 2012). Remarkably, gp120-DC-SIGN interaction triggers a signalling cascade that involves LSP1 and leads to Rho GTPase (a regulator of focal adhesion dynamics) activation (Anand et al., 2009; Prasad et al., 2017; Prasad et al., 2012). This study, thus, support our findings highlighting the key role of LSP1 in the regulation of focal adhesion dynamics and cell migration.

Collectively, our findings provide novel evidence about LSP1 function and its ability to regulate two crucial aspects of cell migration in co-operation with myosin1e: a) actin cytoskeleton assembly (required for lamellipodia formation and dynamics) and b) focal adhesion formation and dynamics. Several questions remain to be addressed. For instance, is the function of myosin1e dependent on its interaction with LSP1? Is LSP1 phosphorylation required for its role in cell migration? Has LSP1 other binding partners in addition to myosin1e? Could LSP1 be a target for novel pharmaceutical treatments of HIV infections? The answers to these questions will certainly help to better understand the role of LPS1 and its interaction with myosin1e not only in the regulation of cell migration and adhesion but also in other processes dependent on actin cytoskeleton remodelling.

## Supporting information

legends for supplementary data

## Acknowledgements

We thank Ms. Gülcan Aydin for excellent technical assistance.

