## Supplementary figures and images for "LSP1-myosin1e bi-molecular complex regulates focal adhesion dynamics and cell migration"

### supplementary figure

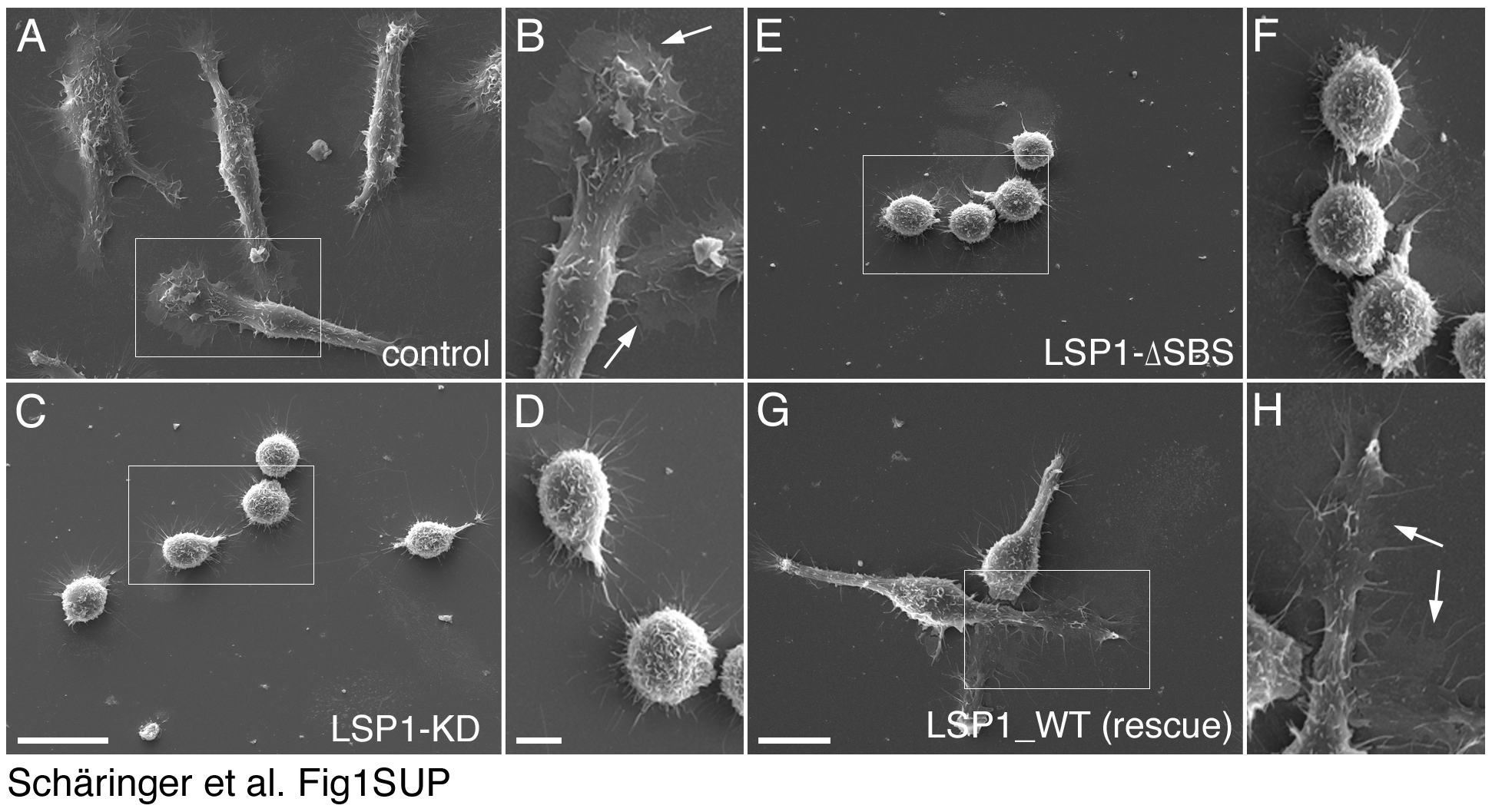
