## Supplementary material for "LSP1-myosin1e bi-molecular complex regulates focal adhesion dynamics and cell migration": legends for supplementary data

Legends for supplementary material

**Figure 1SUP. Analysis of the morphology of J774 cell lines by scanning electron microscopy.** Control J774 cells (A, B), LSP1-deficient cells (C, D), cells expressing the LSP1 mutant unable to interact with myosin1e (LSP1-ΔSBS; E, F) or full-length LSP1 (rescue; G, H) were seeded on glass coverslips and after 24 hours processed for scanning electron microscopy. Both control J774 and J774 cells re-expressing full-length LSP1 acquired an elongated morphology and formed well-developed lamellipodia (arrows in B and H). Conversely, LSP1-deficient cells and cells expressing the LSP1 mutant unable to interact with myosin 1e were round and lacked the large lamellipodia typical of control cells. Scale bars: 5 µm (for B, D, F, H); 10 µm (for G); 20 µm (for A, C, E).

**Movie_Fig3. Focal adhesion dynamics in control and LSP1-deficient J774 macrophages.** Control and LSP1-deficient cells were stably transfected with RFP-zyxin and filmed at 37°C, 5% $CO_2$ in normal culture medium by TIRF microscopy. Note the formation of fewer and smaller focal adhesions in LSP1-deficient cells.

**Movie_Fig5. Impact of LSP1-myosin1e complex on lamellipodia formation and dynamics.** Control, LSP1-deficient J774 macrophages and cells stably expressing the LSP1 mutant ΔSBS were imaged by phase contrast microscopy at 37°C, 5% $CO_2$ in normal culture medium. Control cells formed of a large and unidirectional persistent lamellipodium. By contrast, LSP1-deficient cells and cells expressing the LSP1 mutant ΔSBS formed very small and short-lived lamellipodia.

**Movie_Fig6. Dynamics of LSP1 is affected by its interaction with myosin1e.** Control cells expressing RFP-LSP1, cells expressing the LSP1 mutant ΔSBS (also tagged with RFP) and LSP1-deficient cells expressing full-length LSP1 (RFP tagged; rescue) were filmed at 37°C, 5% $CO_2$ in normal culture medium by TIRF microscopy. Note the more intense LSP1 dynamics in control and rescued cells.

**Movie_Fig7. LSP1-myosin1e is required for efficient focal adhesion formation and dynamics in J774 macrophages.** Cells re-expressing full-length LPS1 (rescue) and cells expressing the LSP1 mutant ΔSBS were stably transfected with turquoise-tagged zyxin and filmed at 37°C, 5% $CO_2$ by TIRF microscopy. Note the formation of fewer and smaller focal adhesions in cells expressing the LSP1 mutant ΔSBS.
